# Hyperactivity Induced By Vapor Inhalation of Nicotine in Male and Female Rats

**DOI:** 10.1101/2024.02.12.579996

**Authors:** Mehrak Javadi-Paydar, Tony M. Kerr, Michael A. Taffe

## Abstract

**Rationale:** Preclinical models of electronic nicotine delivery system (ENDS; “e-cigarette”) use have been rare, so there is an urgent need to develop experimental approaches to evaluate their effects.

**Objective:** To contrast the impact of inhaled nicotine across sex.

**Methods:** Male and female Wistar rats were exposed to vapor from a propylene glycol vehicle (PG), nicotine (NIC; 1-30 mg/mL in PG), or were injected with NIC (0.1-0.8 mg/kg, s.c.), and then assessed for changes in temperature and activity. The antagonist mecamylamine (2 mg/kg) was administered prior to NIC to verify pharmacological specificity. Plasma levels of nicotine and cotinine were determined after inhalation and injection.

**Results:** Activity increased in females for ∼60 minutes after nicotine inhalation, and this was blocked by mecamylamine. A similar magnitude of hyperlocomotion was observed after s.c. administration. Body temperature was reduced after nicotine inhalation by female rats but mecamylamine increased this hypothermia. Increased locomotor activity was observed in male rats if inhalation was extended to 40 minutes or when multiple inhalation epochs were used per session. The temperature of male rats was not altered by nicotine. Plasma nicotine concentrations were slightly lower in male rats than in female rats after 30-minute nicotine vapor inhalation and slightly higher after nicotine injection (1.0 mg/kg, s.c.).

**Conclusions:** Nicotine inhalation increases locomotor activity in male and female rats to a similar or greater extent than by subcutaneous injection. Sex differences were observed, which may be related to lower nicotine plasma levels, lower baseline activity and/or a higher vehicle response in males.

## Introduction

The use of tobacco products represents a major public health concern, particularly when it comes to a resurgence of nicotine use in adolescents via Electronic Nicotine Delivery Systems (ENDS; e-cigarettes; aka “vaping”). The Monitoring the Future survey confirmed high rates of past 30-day vaping in 10^th^ (∼20%) and 12^th^ (∼25%) grade populations in 2019 and 2020 (Johnston et al. 2021). Daily nicotine vaping was reported by about 11.6% of 12^th^ graders in 2019, a demographic where daily cigarette smoking was only 2.4% (Miech et al. 2019). Nicotine exposure differs by sex, since 30-day prevalence of nicotine vaping was 28% (6.9% cigarette use) in male, and 23% (4.0% cigarette use) in female, 12^th^ graders in 2019. Similarly, *daily* cigarette use was reported by 2.8% of male and 1.6% of female 12^th^ graders. Despite the rapid increase in teen vaping from ∼2017-2020, relatively few studies have investigated the impact of nicotine delivered by vapor inhalation in well controlled animal models using ENDS technology until recently. Nicotine may be especially problematic for women since they have less success quitting tobacco use (Perkins 2001; Piper et al. 2010) and nicotine replacement is less effective for smoking cessation in women than men (Perkins et al. 1999; Wetter et al. 1999). Preclinical studies also suggest an enhanced vulnerability to nicotine addiction in female animals (Donny et al. 2000; Faraday et al. 1999; Lynch 2009; Torres et al. 2009), although female rats reach higher plasma and brain nicotine levels compared with male rats after injection of mg/kg adjusted doses (Harrod et al. 2007; Rosecrans 1972), which may contribute to observed sex differences in nicotine-mediated behaviors.

While some initial studies of ENDS based nicotine exposure resorted to injecting e-cigarette vehicle/nicotine preparations into laboratory rodents (LeSage et al. 2016), there has been progress in developing inhalation models for the delivery of several different drugs including cannabinoids (Freels et al. 2020; Javadi-Paydar et al. 2019a; Nguyen et al. 2016b), methamphetamine and other psychomotor stimulants (Marusich et al. 2016; Nguyen et al. 2016a; Nguyen et al. 2017), opioids, including fentanyl (McConnell et al. 2021), heroin (Gutierrez et al. 2021; Gutierrez et al. 2022) and oxycodone (Nguyen et al. 2019). Additional work has focused on developing and deploying rat and mouse models of nicotine vapor inhalation (Cooper et al. 2021; Echeveste Sanchez et al. 2022; Javadi-Paydar et al. 2018; Lallai et al. 2021; Montanari et al. 2020; Roeder et al. 2023; Smith et al. 2020; Zhu et al. 2023). Rodent locomotor behavior is an involuntary response to drugs which can be used as a relatively simple assay of drug potency and efficacy. Although several studies have examined sex differences in the locomotor effects of nicotine when administered by injection to laboratory rodents, relatively few studies have examined nicotine’s effects on locomotion when administered via inhalation. Several different assays of locomotor behavior, including open field assessment by videographic recording (Frie et al. 2023; Gutierrez et al. 2024b; Lallai et al. 2021) or beam break (Alkhlaif and Shelton 2023; Roeder et al. 2023), wheel activity (Gutierrez et al. 2024a; Gutierrez et al. 2024b) and radiotelemetry (Javadi-Paydar et al. 2019a; Javadi-Paydar et al. 2019b), have been used to determine effects of nicotine after vapor inhalation in rats. Despite methodological differences, cotinine levels of ∼20-50 ng/ml of plasma have been consistently reported across laboratory approaches within ∼30 minutes of vapor cessation in rats (Frie et al. 2023; Gutierrez et al. 2024b; Javadi-Paydar et al. 2019b; Lallai et al. 2021; Roeder et al. 2023) and mice (Henderson and Cooper 2021; Lefever et al. 2017). Sometimes the contribution of rat sex has been examined (Frie et al. 2023; Gutierrez et al. 2024b; Lallai et al. 2021) but in other cases not (Alkhlaif and Shelton 2023; Javadi-Paydar et al. 2019a; Javadi-Paydar et al. 2019b; Roeder et al. 2023). In some studies, no locomotor effects of nicotine were observed after the first exposure but a sensitized response appeared after repeated dosing, an effect which may be sex-dependent (Honeycutt et al. 2020; Lallai et al. 2021). Additional work confirmed the locomotor stimulant effect of nicotine inhalation in mice (Echeveste Sanchez et al. 2022; Shao et al. 2019; Zhu et al. 2021), although suppressed activity after nicotine inhalation has also been reported (Lefever et al. 2017). It is very likely that differences in stimulation versus suppression of activity are related to dose and speed of brain entry of nicotine under various dosing methods, which may interact with species, strain or sex.

We have shown that the inhalation of nicotine vapor increases spontaneous locomotor activity and decreases body temperature in male Sprague-Dawley rats, while reductions in activity were observed after nicotine injection (0.4, 0.8 mg/kg, s.c.) (Javadi-Paydar et al. 2019a; Javadi-Paydar et al. 2019b), similar effects have been reported in mice (Honeycutt et al. 2020). Increased activity was found after 15 or 30 minutes of nicotine inhalation (30 mg/mL in the propylene glycol vehicle) in the male rats, lasted only ∼15-30 minutes after vapor exposure and were blocked by pre-inhalation administration of the nicotinic acetylcholine receptor (nAChR) antagonist mecamylamine (Javadi-Paydar et al. 2019b). Plasma nicotine and cotinine levels after 30 minutes of inhalation at the 30 mg/mL vapor concentration were approximately equivalent to plasma levels observed 30 minutes after 0.8 mg/kg, s.c., injection, the dose which reduced activity levels (Javadi-Paydar et al. 2019b). In other studies, it was found that injected nicotine moderately increased open field activity in young adult male and female Sprague-Dawley rats, but suppressed wheel activity, after repeated adolescent exposure to nicotine vapor (Gutierrez et al. 2024b). That study also found that nicotine levels were slightly higher in female versus male Sprague-Dawley rats, and cotinine levels were higher in female Sprague-Dawley and Wistar rats, after nicotine inhalation. A recent study reported that the locomotor activity of female Sprague-Dawley rats in an open field was inhibited after the inhalation of nicotine (24, 59 mg/mL in the vapor vehicle) but increased after nicotine injection (0.4, 1.0 mg/kg, s.c.), with the two routes producing equivalent serum nicotine levels at the high and low dose/concentrations selected (Roeder et al. 2023). Interestingly the same laboratory also reported in a different study that nicotine vapor inhalation increased activity in male rats while not affecting open field activity in female rats (Frie et al. 2023).

This study was designed first to directly compare the impact of nicotine vapor inhalation in male and female rats, with a focus on locomotor activity, since our initial study of telemetry locomotor measures was conducted only in male rats (Javadi-Paydar et al. 2019b) and a follow-up found significant sex differences in open field activity after nicotine injection (Gutierrez et al. 2024b). The second goal was to determine if effects of nicotine inhalation generalize to Wistar rats since our prior behavioral investigations with nicotine vapor have been conducted with Sprague-Dawley rats (Gutierrez et al. 2024b; Javadi-Paydar et al. 2019a; Javadi-Paydar et al. 2019b). This is potentially important since, e.g., we’ve shown that responses to inhaled Δ^9^-tetrahydrocannabinol vary across these common laboratory rat strains (Taffe et al. 2021). More directly, nicotine increased horizontal activity in female, but not male, Long-Evans and Sprague-Dawley rats, with greater effects in Long-Evans females (Faraday et al. 2003).

The third goal was to further determine how variation in time of vapor exposure and drug concentration affect the locomotor effects of nicotine inhalation. Unlike parenteral injection, the methods for vapor inhalation exposure vary across, and within, laboratory and the dose and rate of drug accumulation can be varied by several methodological variations. These may include varying concentration of drug in the vehicle, time of overall inhalation exposure, the number of puffs and volume of vapor delivered per unit time, chamber air evacuation rates, etc. This study determined if changes in drug concentration in the vehicle and time of exposure alter the locomotor responses.

## Methods

### Subjects

Groups of male (N=31 total) and female (N=24 total) Wistar rats (Charles River, New York) were housed in humidity and temperature-controlled (23 ± 1 °C) vivaria on 12:12 h light:dark cycles. Groups 1-3 consisted of subgroups of male and female rats (N=8 per sex; one Group 1 male was lost prior to the initiation of these studies) obtained as cohorts and treated similarly by Group prior to this study, as described below. Group 4 consisted of experimentally naïve male rats. All experimental procedures took place during the animals’ dark (active) cycle. Animals had ad libitum access to food and water in their home cages. All procedures were conducted under protocols approved by the Institutional Care and Use Committee of The Scripps Research Institute and in a manner consistent with the Guide for the Care and Use of Laboratory Animals (National Research Council (U.S.) Committee for the Update of the Guide for the Care and Use of Laboratory Animals. et al., 2011).

### Drugs

(−)-Nicotine bitartrate and mecamylamine hydrochloride (Sigma-Aldrich) were dissolved in 0.9% sterile saline for subcutaneous (s.c.) injection. The doses of nicotine (0.1, 0.2, 0.4 and 0.8 mg/kg) and mecamylamine (2.0 mg/kg) are expressed as the bitartrate and HCl, respectively. D-methamphetamine HCl (NIDA Drug Supply) was dissolved in 0.9% sterile saline to a concentration of 1.0 mg/ml of solution for s.c. acute challenges. A constant injection volume of 1 ml/kg was used for all drugs. The inhalation exposure was to nicotine (NIC; 1, 5, 10, 20 and 30 mg/mL; concentrations are expressed as the bitartrate) in the propylene glycol (PG) vehicle. Four 10-s vapor puffs were delivered with 2-s intervals every 5 minutes, which resulted in use of approximately 0.125 ml in a 40 minute exposure session as in prior studies (Nguyen et al. 2016b). The vacuum was turned off for the 4-minute, 12-second interval between vapor deliveries and then turned up to ∼3-5 L/minutes at the conclusion of sessions for ∼5 minutes to facilitate complete chamber clearance for subject removal.

### Procedures

#### Radiotransmitter implantation

Rats were anesthetized with an isoflurane/oxygen vapor mixture (isoflurane 5% induction, 3% maintenance), and sterile radiotelemetry transmitters (Data Sciences International; TA-F40) were implanted in the abdominal cavity through an incision along the abdominal midline posterior to the xyphoid space as previously described (Aarde et al. 2015; Taffe et al. 2015). Absorbable sutures were used to close the abdominal muscle incision and the skin incision was closed with the tissue adhesive (3M Vetbond Tissue Adhesive; 3M, St Paul, MN). A minimum of 7 days was allowed for surgical recovery prior to starting experiments. For the first three days of the recovery period, an antibiotic Cefazolin (0.4 g/ml, 2.0 ml/kg, s.c.) and an analgesic, flunixin (2.5 mg/ml, 2.0 ml/kg, s.c.) daily for 3 days.

#### Radiotelemetry recording

Measures of locomotor activity and body temperature were collected every 5 minutes while animals were housed in the inhalation chambers in a dark testing room (dim red-light illumination), separate from the vivarium, during the (vivarium) dark cycle. Radiotelemetry transmissions were collected via receiver plates (Data Sciences International; RPC-1) placed under the cages as described in prior investigations (Javadi-Paydar et al. 2018; Taffe et al. 2015). The ambient temperature for the studies was 22 ± 1 °C. Sessions started with a 30-min interval in the recording cage to determine a pre-treatment baseline of activity and temperature. Animals were then exposed to the experimental vapor inhalation conditions while recording continued. In some cases, rats were briefly removed from the chambers for injection with experimental pre-treatments. The three telemetry observations obtained prior to drug treatment were averaged as the baseline for analysis, and timepoints are referenced to the start of vapor exposure.

#### Vapor Inhalation

Nicotine was delivered by vapor inhalation as previously described (Gutierrez et al. 2024b; Javadi-Paydar et al. 2019b). In brief, an e-vape controller (Model SSV-1; 3.3 volts; La Jolla Alcohol Research, Inc, La Jolla, CA, USA) was used to deliver a scheduled series of puffs from e-cigarette cartridges (Protank 3 Atomizer, MT32 coil operating at 2.2 ohms, by Kanger Tech; Shenzhen Kanger Technology Co.,LTD; Fuyong Town, Shenzhen, China) under computerized control (Control Cube 1; La Jolla Alcohol Research, Inc, La Jolla, CA, USA). The chamber (259 mm X 234 mm X 209 mm Allentown, Inc. Allentown, NJ) air was vacuum controlled by a chamber exhaust valve (i.e., a “pull” system) to flow room ambient air through an intake valve at ∼1 L per minute. Drug dosing conditions were manipulated by altering the concentration in the PG vehicle.

### Temperature and Activity Experiments

#### *Experiment 1*. Behavioral response in male and female rats **(Group 1)**

The Group 1 male (N = 7; 50 weeks of age, mean bodyweight 678.0 SEM 13.11 g) and female (N = 8, 50 weeks of age, mean bodyweight 344.4 SEM 13.34 g) rats were exposed to inhalation of propylene glycol (PG) vapor versus nicotine (30 mg/mL in PG) for 30 min to determine any sex differences with the approach used in our prior study (Javadi-Paydar et al. 2019b). These rats were initially used for experiments described previously (Javadi-Paydar et al. 2018) involving inhalation of THC (12.5-200 mg/mL) and CBD (100 and 400 mg/mL) vapor, no more frequently than once per week, for ∼7 months.

#### *Experiment 2*. Nicotine vapor duration-response in male rats **(Group 1)**

The male rats were next exposed to inhalation of PG vapor versus nicotine (30 mg/mL) for 15, 30 and 40 minutes to determine if the lack of locomotor effect in Experiment 1 was likely due to an insufficient nicotine dose.

#### *Experiment 3*. Nicotine vapor concentration-response in female rats **(Group 2)**

To determine the range of effective inhalation conditions in female rats, the Group 2 female rats (N = 8; 15 weeks of age, mean bodyweight 194.0 SEM 2.94 g) were exposed to inhalation of propylene glycol (PG) vapor versus nicotine (10, 20 and 30 mg/mL) vapor for 15 minutes with dose conditions evaluated in a counter-balanced order. Then they were exposed to inhalation of PG vapor versus lower concentrations of nicotine (1, 5 and 10 mg/mL) vapor for 15 minutes, again with these four dose conditions evaluated in a counter-balanced order. These rats were initially used for other experiments involving exposure to THC and nicotine vapor inhalation, no more frequently than once per week, for ∼2 months.

#### *Experiment 4*. Effect of mecamylamine on the response to inhalation of nicotine **(Group 2)**

Our prior study in male Sprague-Dawley rats found that locomotor effects of inhaled nicotine were most robust when rats were exposed to multiple vapor inhalation intervals per session (Javadi-Paydar et al. 2019b), due to increased activity in response to the first puffs of the session regardless of the presence of nicotine. This experiment used the noncompetitive nAChR antagonist mecamylamine (Young et al. 2001) to determine the pharmacological specificity of the locomotor response to two nicotine vapor intervals per session.

The Group 2 female rats were injected with saline or mecamylamine (2 mg/kg in saline, s.c.) 15 min prior to inhalation of PG or nicotine (30 mg/mL) vapor for 15 minutes, followed by a second 15-minute interval of PG or nicotine (30 mg/mL) inhalation two hours later. Treatment conditions (two pre-treatment injections, two initial vapor conditions, two final vapor conditions) were counterbalanced in order across the group, and a total of 8 sessions were completed per rat.

The Group 2 male rats (N = 8; 23 weeks of age, mean bodyweight 575.4 SEM 15.20 g) were likewise exposed to 15 min inhalation of PG or nicotine vapor (30 mg/mL), followed with a second PG or nicotine inhalation session 2 hours later, with all four conditions tested in a counter balanced order. They were then evaluated for the effects of saline or mecamylamine (2 mg/kg in saline, s.c.) in a counterbalanced order. In this latter study, injections were 15 minutes prior to inhalation of PG for 15 min followed one hour later with PG or nicotine inhalation (30 mg/mL) for 30 minutes. These rats were initially used for other experiments involving exposure to THC and nicotine vapor inhalation, no more frequently than once per week, for ∼2 months.

#### *Experiment 5*. Subcutaneous nicotine dose-response in adult male and female rats **(Group 2)**

Female Wistar rats (Group 2; N = 8; 38 weeks of age, mean bodyweight 367.6 SEM 12.90 g at the start of this study) were injected with saline or nicotine (0.1, 0.2, 0.4 and 0.8 mg/kg, s.c.) in a counterbalanced order. They were next injected with saline or mecamylamine (2 mg/kg, s.c.) 15 min prior to injection of saline or nicotine (0.8 mg/kg, s.c.). Doses were selected based on prior and ongoing locomotor studies in the laboratory with compounds with a similar range of in vivo pharmacological effects.

Male Wistar rats (Group 2; N = 8; 38 weeks of age, mean bodyweight 699.4 SEM 26.88 g at the start of this study) were injected with saline or nicotine (0.1, 0.2, 0.4 and 0.8 mg/kg, s.c.). Because the locomotor response to nicotine was of small magnitude, following this, they were tested with injection of 1.0 mg/kg, s.c. of methamphetamine, or saline, in a counterbalanced order, as a positive control to determine if they would express a locomotor response to pharmacological challenge under these conditions.

#### Plasma collection after vapor inhalation and subcutaneous injection **(Group 3, 4)**

To assess plasma nicotine and cotinine, blood samples (∼0.5 ml) were withdrawn from the jugular vein under inhalation anesthesia from **<u>Group 3</u>** male (N = 8; 32 weeks of age, mean bodyweight 602.38 SEM 19.44 g) and female (N = 8; 32 weeks of age, mean bodyweight 319.88 SEM 8.98 g) Wistar rats immediately after inhalation of nicotine (30 mg/mL, 30 minutes) vapor and then 90 minutes later. Samples were also obtained from this group two weeks later, 15 and 90 minutes after nicotine injection (1.0 mg/kg, s.c.); the 14 day interval was interposed between blood sampling experiments to align with guidelines for blood recovery following multiple samples (Diehl et al. 2001). These rats had initially been used for other plasma sampling experiments following inhalation of THC (50-200 mg/ml in PG) vapor, following twice daily THC inhalation during postnatal days 36-40 as described previously (Nguyen et al. 2018).

Male (**<u>Group 4</u>**; N = 8; 37 weeks of age, mean bodyweight 676.1 SEM 25.35 g) Wistar rats were used for sampling blood immediately after nicotine injection (0.2, 0.4, 0.8 mg/kg, s.c.) as well as after inhalation of nicotine vapor (10 and 30 mg/mL for 30 minutes). Blood samples were withdrawn 15 minutes after the injection and 35 minutes after the start of vapor inhalation. No other experiments were performed with Group 4 prior to the current experiment and a minimum 7 day interval was interposed between blood sampling experiments to align with guidelines for blood recovery following single samples (Diehl et al. 2001). One rat was euthanized due to developing a solid mass prior to the inhalation experiment, thus N=7 for that study.

### Nicotine and Cotinine Analysis

Plasma nicotine and cotinine content was quantified using liquid chromatography/mass spectrometry (LCMS) as previously described (Javadi-Paydar et al. 2019b). Briefly, 50 µl of plasma were mixed with 50 µl of deuterated internal standard (100 ng/ml cotinine-d3 and nicotine-d4; Cerilliant). Nicotine and cotinine (and the internal standards) were extracted into 900 µl of acetonitrile and then dried. Samples were reconstituted in 100 µL of an acetonitrile/water (9:1) mixture. Separation was performed on an Agilent LC1200 with an Agilent Poroshell 120 HILIC column (2.1mm x 100mm; 2.7 um) using an isocratic mobile phase composition of acetonitrile/water (90:10) with 0.2% formic acid at a flow rate of 325 µL/min. Nicotine and cotinine were quantified using an Agilent MSD6130 single quadrupole interfaced with electrospray ionization and selected ion monitoring [nicotine (m/z=163.1), nicotine-d4 (m/z=167.1), cotinine (m/z=177.1) and cotinine-d3 (m/z=180.1)]. Calibration curves were generated daily using a concentration range of 0-200 ng/mL with observed correlation coefficients of 0.999.

### Data Analysis

The body temperature (°C) and activity rate (counts per minute) were collected via the radio-telemetry system on a 5-minute schedule and analyzed in 30-minute averages (the time point refers to the ending time, i.e. 60 = average of 35-60 minute samples). Any missing temperature values (e.g., due to radio interference or animal’s location within the chamber at the time of sampling) were interpolated from preceding and succeeding values. Telemetry data were analyzed with Analysis of Variance (ANOVA) with repeated measures factors for the Drug Treatment Condition and the Time after initiation of vapor (or post-injection). Any significant effects in the main analysis were followed with post-hoc analysis using Tukey correction for all multi-level comparisons and Sidak for two-level comparisons. The analysis of plasma nicotine and cotinine concentrations for Group 3 used a third factor for Sex, and the post hoc analysis was limited to orthogonal comparisons (cells that differ only by one factor) using Bonferroni correction. All analysis used Prism for Windows (GraphPad Software, Inc, San Diego CA).

## Results

### Effects of vapor inhalation of nicotine

Inhalation of nicotine vapor (30 mg/ml concentration) for 30 minutes altered the activity and body temperature of the Group 1 female, but not Group 1 male, rats (**Figure 1**).

**Figure 1.**
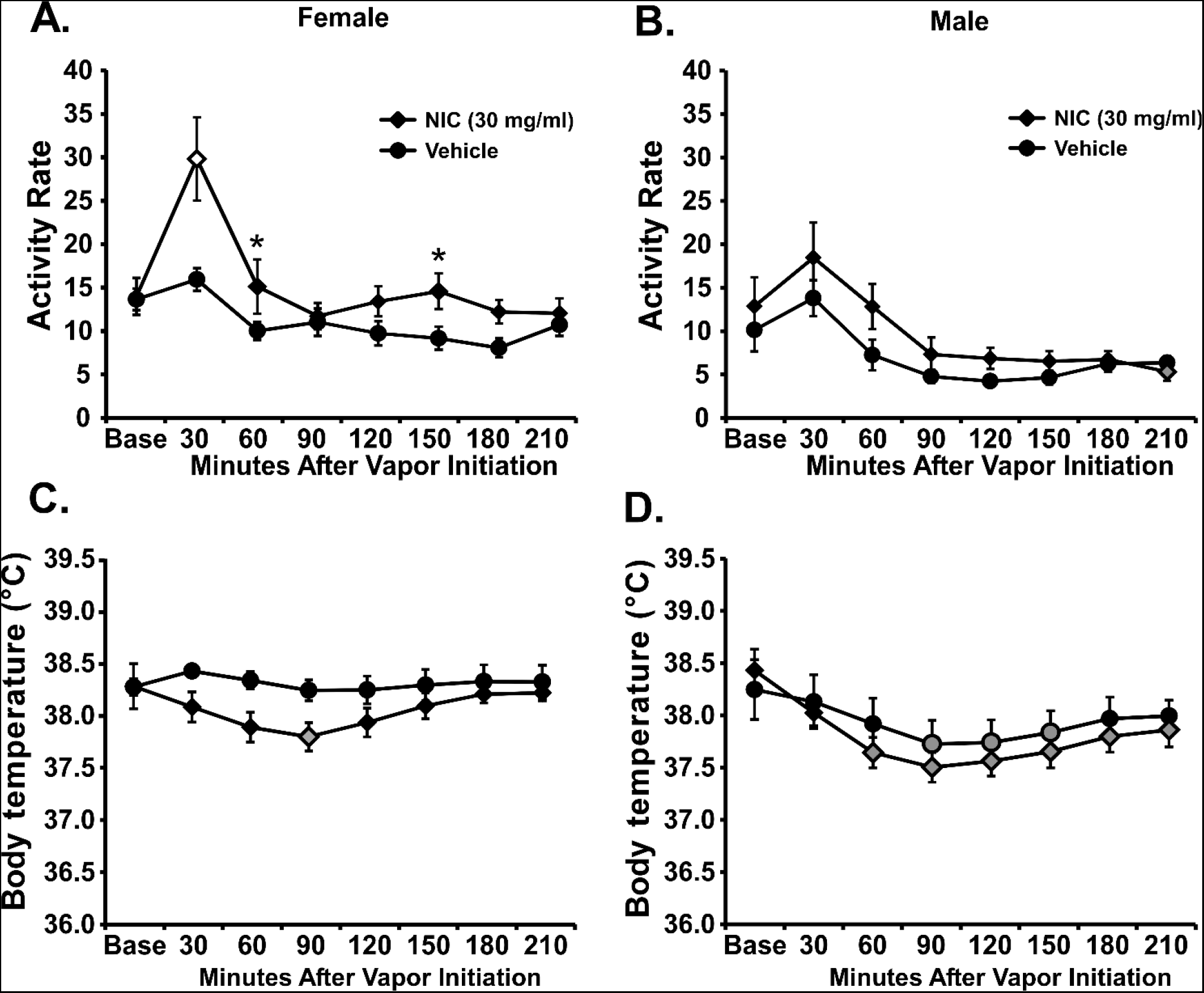
Mean (±SEM for N=7 male; N=8 female) A, B) activity rate and C, D) temperature after inhalation of the PG vehicle or nicotine (NIC; 30 mg/mL; 30 min). Open symbols indicate a significant difference from both the PG and the within-treatment baseline at a given time-point and shaded symbols indicate a significant difference from the baseline only. A significant difference from PG inhalation conditions is indicated by ^*^.

***Female rats* (Group 1)** The ANOVA confirmed significant effects of Time Post-initiation [F (7, 49) = 21.85; P < 0.0001], of Vapor Condition [F (1, 7) = 6.276; P < 0.05], and of the interaction of factors [F (7, 49) = 2.93; P < 0.05] on the activity of female rats. The post-hoc test further confirmed that activity was significantly increased compared with the baseline after nicotine inhalation, but not after PG inhalation. The post-hoc test also confirmed significant differences in activity 30, 60 and 150 minutes after initiation of nicotine vapor compared with the PG control condition (**Figure 1A**). Analysis of body temperature confirmed a significant effect of Time Post-initiation [F (7, 49) = 3.021; P < 0.05], but not of Vapor Condition or of the interaction of factors. However, the post-hoc test confirmed that body temperature was significantly decreased compared with the baseline after inhalation of nicotine, but not after PG inhalation (**Figure 1C**).

***Male rats* (Group 1)** Vapor inhalation of nicotine (30 mg/ml) for 30 minutes did not affect the temperature or activity of the older male rats to the same extent. While the ANOVA confirmed a significant effect of Time Post-initiation [F (7, 42) = 8.76; P < 0.0001], this did not differ significantly across the two vapor inhalation conditions (**Figure 1B**), although the post-hoc confirmed that activity was lower baseline at the final timepoint after nicotine inhalation. Likewise, there was no effect of vapor inhalation condition on body temperature. The ANOVA confirmed a significant effect of Time Post-initiation [F (7, 42) = 16.68; P < 0.0001], but not of the vapor drug condition or of the interaction of factors (**Figure 1D**). The post-hoc test confirmed that temperature was lower than baseline after PG (90-150 minutes) or nicotine (60-210 minutes) inhalation.

To further explore the lack of nicotine effect in these males, they were exposed to the 30 mg/mL concentration of nicotine vapor for varying durations of inhalation (15, 30, 40 min) in Experiment 2. The ANOVA confirmed significant effects of Time Post-initiation [F (7, 42) = 13.47; P < 0.0001], and the interaction of Time with Vapor Duration [F (21, 126) = 1.85; P < 0.05] on activity rates. The post-hoc test confirmed that activity was significantly increased compared with the baseline after inhalation of nicotine for 40 minutes and significantly higher compared with PG inhalation 30 and 60 minutes after the start of the 30- and 40-minute inhalation conditions (**Figure 2A**).

**Figure 2.**
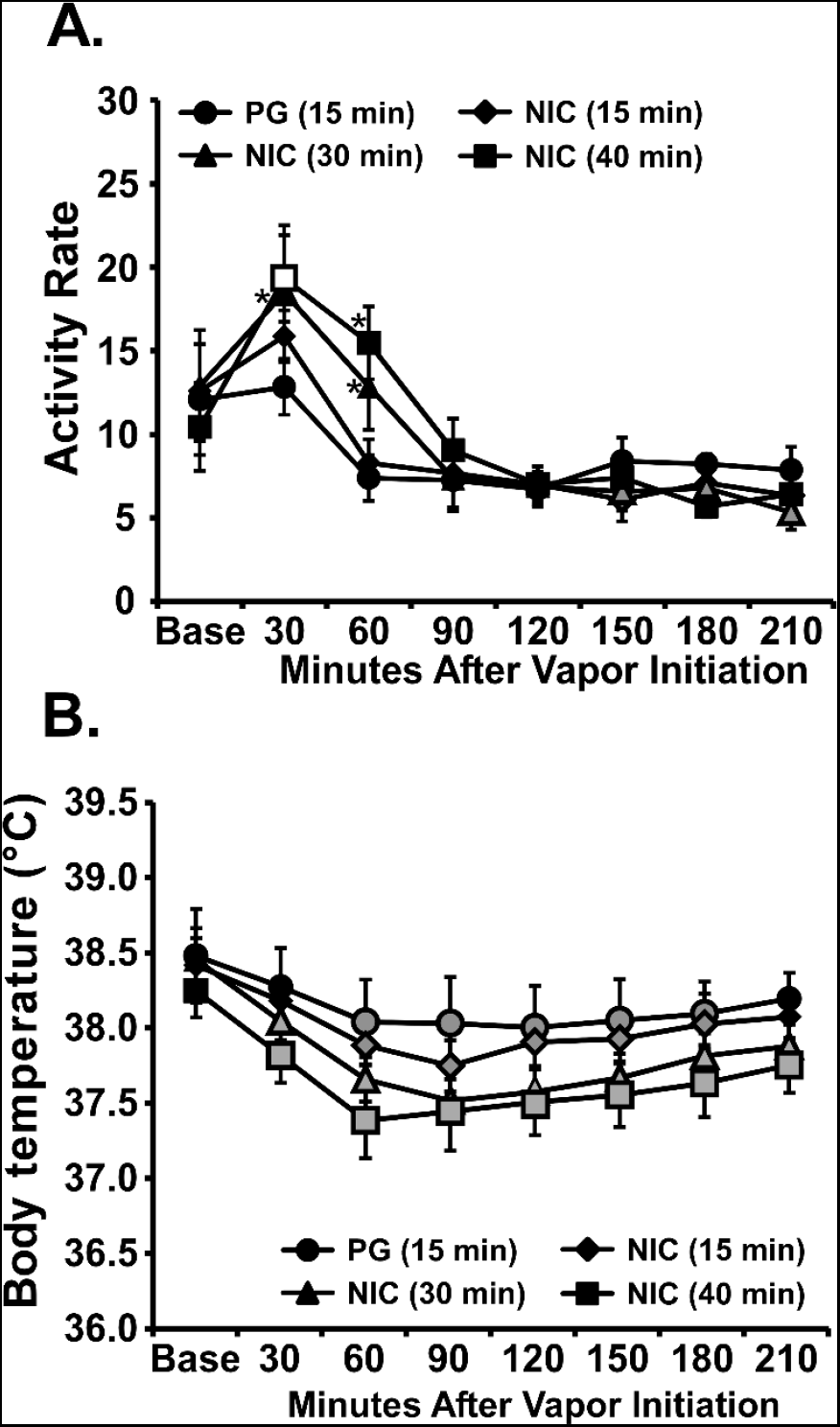
Mean (±SEM; N=7 male) A) Activity rate and B) temperature after inhalation of the PG vehicle or nicotine (NIC; 30 mg/mL; 15, 30, 40 min). Open symbols indicate a significant difference from both the within-treatment baseline and the vehicle at a given time-point and shaded symbols indicate a significant difference from the baseline only. A significant difference from the PG inhalation condition is indicated by ^*^.

The temperature of male rats was decreased by NIC inhalation in a duration-dependent manner **(Fig. 2B)** and the ANOVA confirmed a significant effect of Time Post-initiation [F (7, 42) = 11.55; P < 0.0001]. Post-hoc exploration confirmed that the body temperature was significantly different from both the baseline and PG inhalation condition after 30- and 40-minute nicotine inhalation.

### Plasma nicotine and cotinine after vapor inhalation

Plasma nicotine and cotinine levels were assessed to determine the extent to which sex differences in locomotor response might be due to differential nicotine exposure or metabolism.

This study confirmed a small sex difference in plasma nicotine after identical inhalation conditions **(Figure 3)** in the **<u>Group 3</u>** rats. The three-way ANOVA confirmed significant effects of Analyte [F (1, 14) = 116.5; P<0.0001], of Time [F (1, 14) = 29.51; P<0.0001], of the interaction of Analyte and Time [F (1, 14) = 123.9; P<0.0001] and the interaction of all three factors [F (1, 14) = 8.50; P=0.05] on plasma concentration after inhalation **(Figure 3A)**. The post-hoc test confirmed that plasma nicotine was lower for each sex, and cotinine levels were higher in the females, at the second timepoint. In addition, plasma cotinine concentrations were higher than nicotine at the second time-point. Finally, male nicotine concentrations were lower than females at the post-session timepoint. The three-way ANOVA confirmed significant effects of Sex [F (1, 56) = 14.29; P<0.0005] and of the interaction of Analyte and Time [F (1, 56) = 276.2; P<0.0001] on plasma concentration after injection **(Figure 3B)**. The post-hoc test confirmed that plasma nicotine concentrations were lower, and cotinine concentrations were higher, at the second timepoint for each sex. Nicotine and cotinine levels were also significantly different at each timepoint for each sex. Finally, nicotine concentrations were higher in the male rats compared with the female rats 15 minutes after injection.

**Figure 3.**
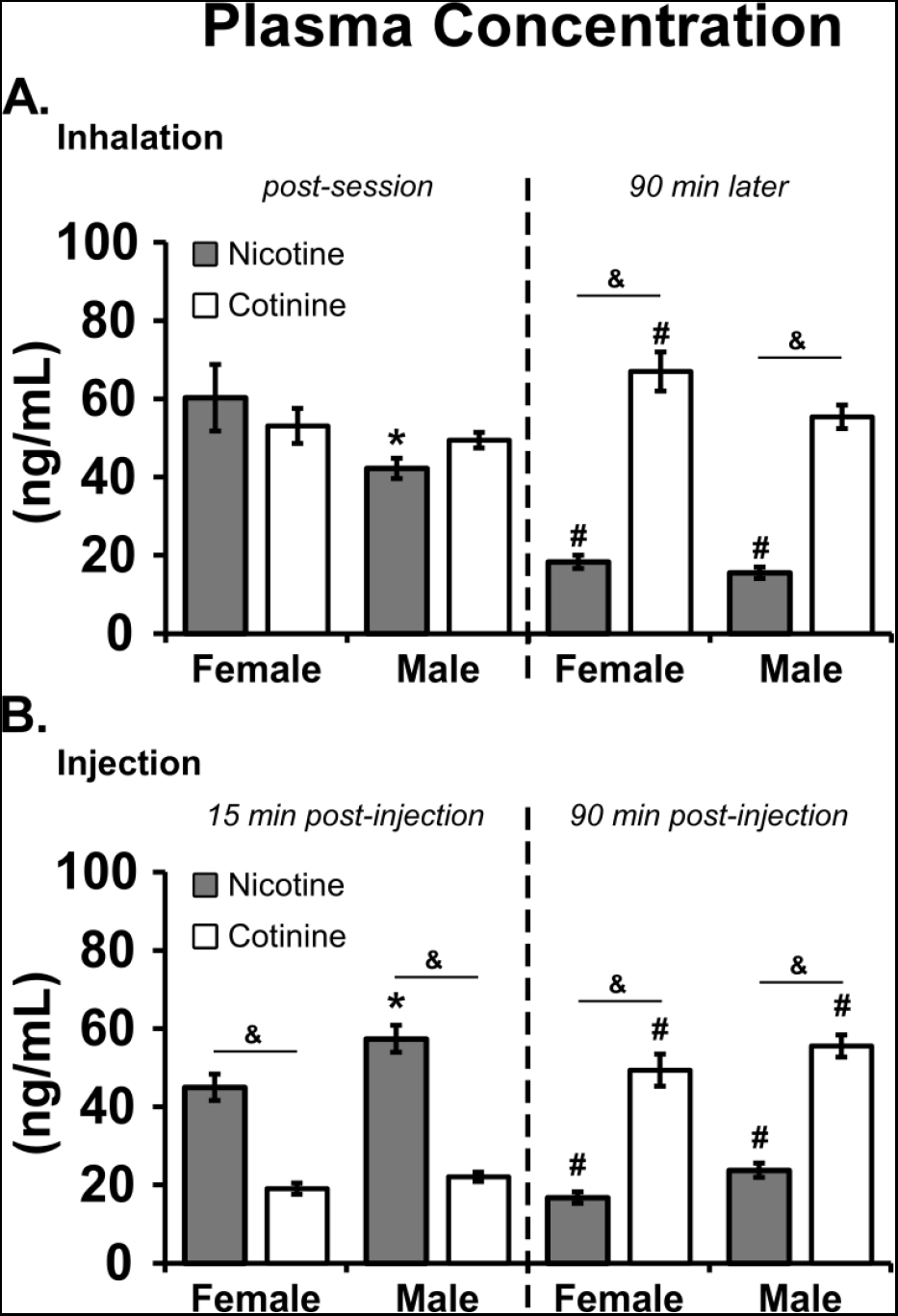
Mean (N=8; ±SEM) plasma nicotine and cotinine after A) inhalation of nicotine (30 mg/mL) vapor for 30 minutes or B) nicotine injection (1.0 mg/kg, s.c.) in age matched (30 weeks) male and female Wistar rats. ^*^=significant sex difference within time and analyte; #=significant difference across time within analyte and sex; &=significant analyte difference.

### Concentration-response for nicotine vapor

In Experiment 3, the activity of the **<u>Group 2</u>** female rats was increased by 15 minutes of inhalation of nicotine vapor in a dose-dependent manner in both high (10-30 mg/ml; **Figure 4A**) and low (1-10 mg/ml; **Figure 4B**) concentration experiments. Analysis was limited to the first 180 minutes for the high concentration (10-30 mg/ml) experiment because data were inadvertently unavailable for the final timepoint for two individuals in all four conditions; data were also unavailable for two animals for the 120-210 minute intervals in the PG inhalation condition, thus mixed-effects analysis was used. The analysis confirmed significant effects of Time Post-initiation [F (6, 42) = 21.24; P < 0.0001], and of Vapor Condition [F (3, 21) = 5.574; P < 0.01], and the interaction of factors [F (18, 120) = 1.70; P < 0.05] on activity (**Figure 4A**). The post-hoc analysis confirmed that activity was significantly increased compared with the baseline 30 minutes after inhalation of 10 and 30 mg/mL nicotine, and *decreased* compared with the baseline 60 minutes after the PG exposure. The post-hoc test also confirmed significant increases in activity compared with PG inhalation for the 10 mg/mL (30 minutes after vapor initiation), or 20 and 30 mg/mL (30 and 60 minutes) nicotine conditions. In the lower concentration (1-10 mg/mL) experiment, the ANOVA confirmed significant effects of Time Post-initiation [F (7, 49) = 21.47; P<0.0001], but not of Vapor Conditions or by the interaction of factors. Activity was significantly increased compared with the baseline after inhalation of 1-10 mg/mL (30 minutes) nicotine, but not after the PG condition. The post-hoc test also confirmed significant differences from PG 30 minutes after the after inhalation of 5-10 mg/mL (30 minutes) nicotine vapor. The *temperature* of female rats was significantly altered by Time Post-injection, but not by Dose Condition nor by the interaction of factors, in both the 10-30 mg/mL [F (6, 42) = 10.46; P<0.0001] and 1-10 mg/mL [F (7, 49) = 2.71; P<0.05] concentration experiments (**Figure 4C, D**).

**Figure 4.**
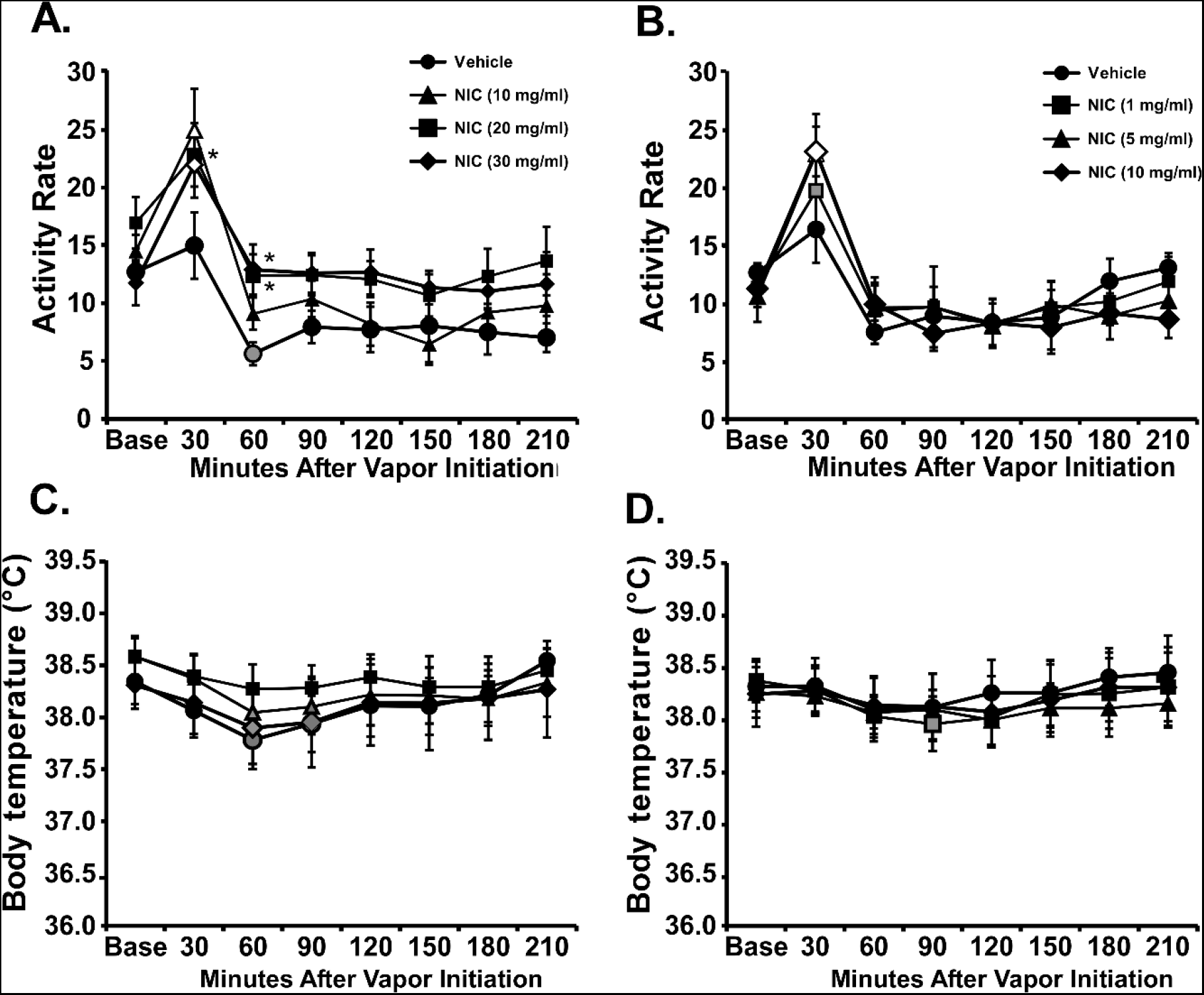
Mean (±SEM; N=8 female) A, B) activity rate and C, D) temperature after inhalation of the PG vehicle or nicotine (A, C; NIC 10, 20, 30 mg/mL; B, D; NIC 1, 5, 10 mg/ml; 15 min). Open symbols indicate a significant difference from both the within-treatment baseline and the vehicle at a given time-point, while shaded symbols indicate a significant difference from the baseline only. A significant difference from PG inhalation condition is indicated by ^*^.

### Effect of mecamylamine on the response to nicotine vapor inhalation

The activity of the **<u>Group 2 female rats</u>** was increased by NIC inhalation, an effect that was attenuated by mecamylamine, in Experiment 4 (**Figure 5**). The statistical analysis was conducted on groups of four treatment sessions, differentiated by the presence or absence of nicotine on the first exposure of the session. The analysis of sessions in which nicotine vapor was delivered first (**Figure 5A**) confirmed significant effects of Time Post-initiation [F (7, 49) = 49.51; P<0.0001], of Treatment Condition [F (3, 21) = 6.61; P<0.01] and of the interaction of Time and Treatment Condition [F (21, 147) = 2.93; P<0.0001]. The Tukey post-hoc test confirmed that an increase in activity after the first nicotine inhalation was significantly reduced by mecamylamine pre-treatment (**Figure 5A**).

**Figure 5.**
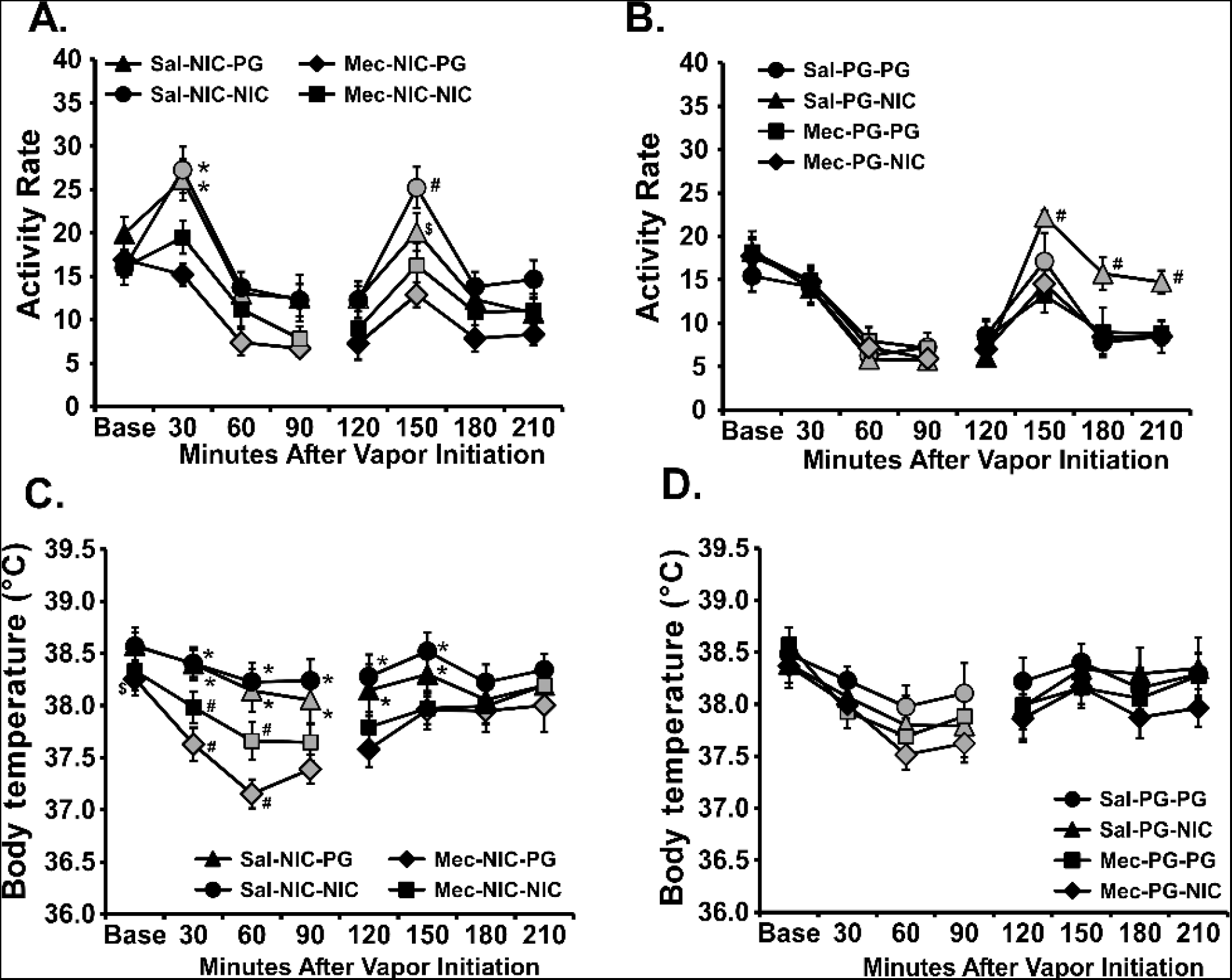
Mean (±SEM; N=8 female) A, B) activity rate and C, D) temperature after inhalation of the PG vehicle or nicotine (NIC; 30 mg/mL; 15 min) twice per session, with Saline or Mecamylamine pre-treatment. All eight treatment conditions were counterbalanced in one experiment and are presented by the initial vapor condition for clarity. Shaded symbols indicate a significant difference from the baseline or 120-minute timepoint, within treatment condition. For a given time-point, a significant difference from both mecamylamine pre-treatment conditions is indicated by ^*^, a difference from all other conditions with # and a difference between Saline and Mecamylamine pre-treatments for the same vapor sequence with $.

Activity was likewise increased more after the second inhalation conditions where nicotine was available compared with the respective PG inhalation and this was attenuated when mecamylamine had been administered before the session.

For the sessions in which PG vapor was delivered first in the session (**Figure 5B**), the ANOVA confirmed significant effects of Time Post-initiation [F (7, 49 = 38.94; P<0.0001], and of the interaction of Time and vapor conditions [F (21, 147) = 3.28; P<0.0001]. The Tukey post hoc test confirmed first that activity was reduced compared with the baseline after the initial PG inhalation in all four conditions. Activity was also significantly increased, relative to the 120-minute timepoint, in all three conditions other than the PG-PG after mecamylamine. However, activity was higher in the saline-nicotine-nicotine condition compared with all three other conditions from 150-210 minutes after the start of the first inhalation.

Body temperature was lowered by mecamylamine injection and nicotine inhalation, alone and in combination, when the nicotine was the first exposure of the session (**Figure 5C**). The ANOVA confirmed significant effects of Time Post-initiation [F (7, 49) = 21.74; P<0.0001], of Treatment Condition [F (3, 21) = 4.85; P=0.01] and of the interaction of Time and Treatment Condition [F (21, 147) = 3.264; P<0.0001]. The post-hoc test confirmed that body temperature was significantly different from the baseline in the saline-nicotine-PG (60, 90 minutes), mecamylamine-nicotine-PG (30, 60, 90 minutes) and mecamylamine-nicotine-nicotine (60, 90 minutes) inhalation conditions. Furthermore, temperature was significantly lower after mecamylamine injection compared to saline injection from 30-150 minutes after the start of the initial inhalation.

Body temperature was lower after the start of inhalation when PG was the first condition of the session, but this was not affected significantly by vapor inhalation (**Figure 5D**). The ANOVA confirmed significant effects of Time Post-Initiation [F (7, 49) = 22.15; P<0.0001]; but not of Treatment Condition or of the interaction of Time and Treatment Condition. The Tukey post-hoc test confirmed that body temperature was significantly different from the first baseline 60-90 minutes after vapor initiation in all conditions and at the 30-minute time point in the Mecamylamine-PG-PG condition. Temperature was also lower in the mecamylamine-PG-nicotine condition compared with the saline-PG-PG condition 60-90 minutes after the start of inhalation.

The activity of the **<u>Group 2 male rats</u>** was first evaluated in four counterbalanced conditions in which they were exposed to PG or nicotine (30 mg/mL) vapor for 15 minutes twice, with a 2-hour interval between exposures (**Figure 6A**). The ANOVA confirmed significant effects of Time Post-initiation [F (3, 21) = 8.196; P<0.01], and of the interaction of Time and Vapor condition [F (9, 63) = 2.554; P<0.05] on activity. The post-hoc test confirmed that activity was significantly increased over baseline 30 minutes after initiation of the first inhalation session in all conditions. Activity was also significantly higher in the nicotine-nicotine condition compared with nicotine-PG at this timepoint. The post-hoc test further confirmed that activity was only increased in the 150-minute timepoint compared with the 120-minute timepoint in the PG-nicotine and nicotine-nicotine conditions. Likewise, activity in the nicotine-nicotine condition was higher than after either PG-PG or nicotine-PG conditions and higher in the PG-nicotine condition compared with the PG-PG condition. No significant effects of vapor inhalation on body temperature were observed in this Experiment (**Figure 6B**).

**Figure 6.**
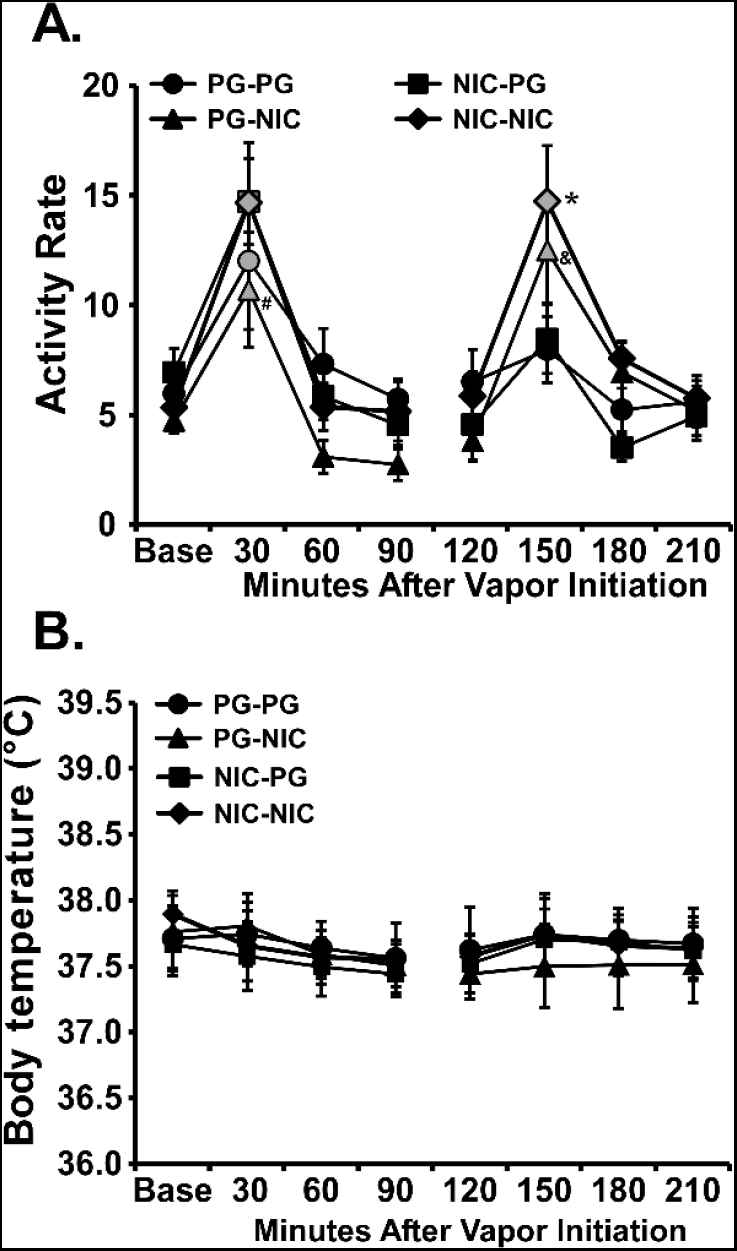
Mean (±SEM; N=8 male) A) activity rate and B) temperature after inhalation of the PG vehicle or nicotine (NIC; 30 mg/mL; 15 min) twice per session. Shaded symbols indicate a significant difference from the baseline or 120-minute timepoint, within treatment condition. For a given time-point, a significant difference from both PG inhalation conditions is indicated by ^*^, a significant difference between PG-NIC and PG-PG for sessions with &, and between PG-NIC and NIC-NIC and sessions with #.

In the mecamylamine experiment, the ANOVA confirmed significant effects of Time Post-injection [F (8, 56) = 6.21; P<0.0001], and of the interaction of Time and Treatment Condition [F (24, 168) = 2.17; P<0.005] on activity of the male rats (**Figure 7**). The post-hoc test first confirmed that activity rate was suppressed during the second vapor exposure compared with the pre-study baseline after Saline injection but not after mecamylamine injection. Thus, the comparison of the effect of the 30-minute inhalation conditions is benchmarked to the 60-minute post-injection time. The post-hoc test confirmed that activity rates increased relative to this level in the 30-minute interval following vapor exposure in all conditions except the mecamylamine-PG condition and remained elevated in the Saline-nicotine condition through 150 minutes after the initial injection. In addition, activity was significantly higher in the Saline-nicotine compared with the mecamylamine-nicotine condition in the 180-minute time bin and higher in the Saline-nicotine compared with the Saline-PG conditions in the 120-minute time bin. Body temperature was altered by mecamylamine administered prior to NIC inhalation conditions (data not shown). The ANOVA confirmed significant effects of Time Post-initiation [F (8, 56) = 6.66; P<0.0001] and of the interaction of Time and Treatment Condition [F (24, 168) = 5.57; P<0.0001] on temperature. The post-hoc test confirmed that body temperature was significantly lower compared with the baseline in the saline-nicotine (120 minutes), mecamylamine-PG (60-150 minutes), and mecamylamine-nicotine (60-150 minutes) conditions. Temperature was also lower after mecamylamine-PG compared with saline-PG (90-150 minutes) and after the mecamylamine-nicotine compared with saline-nicotine (60-150 minutes) conditions.

**Figure 7:**
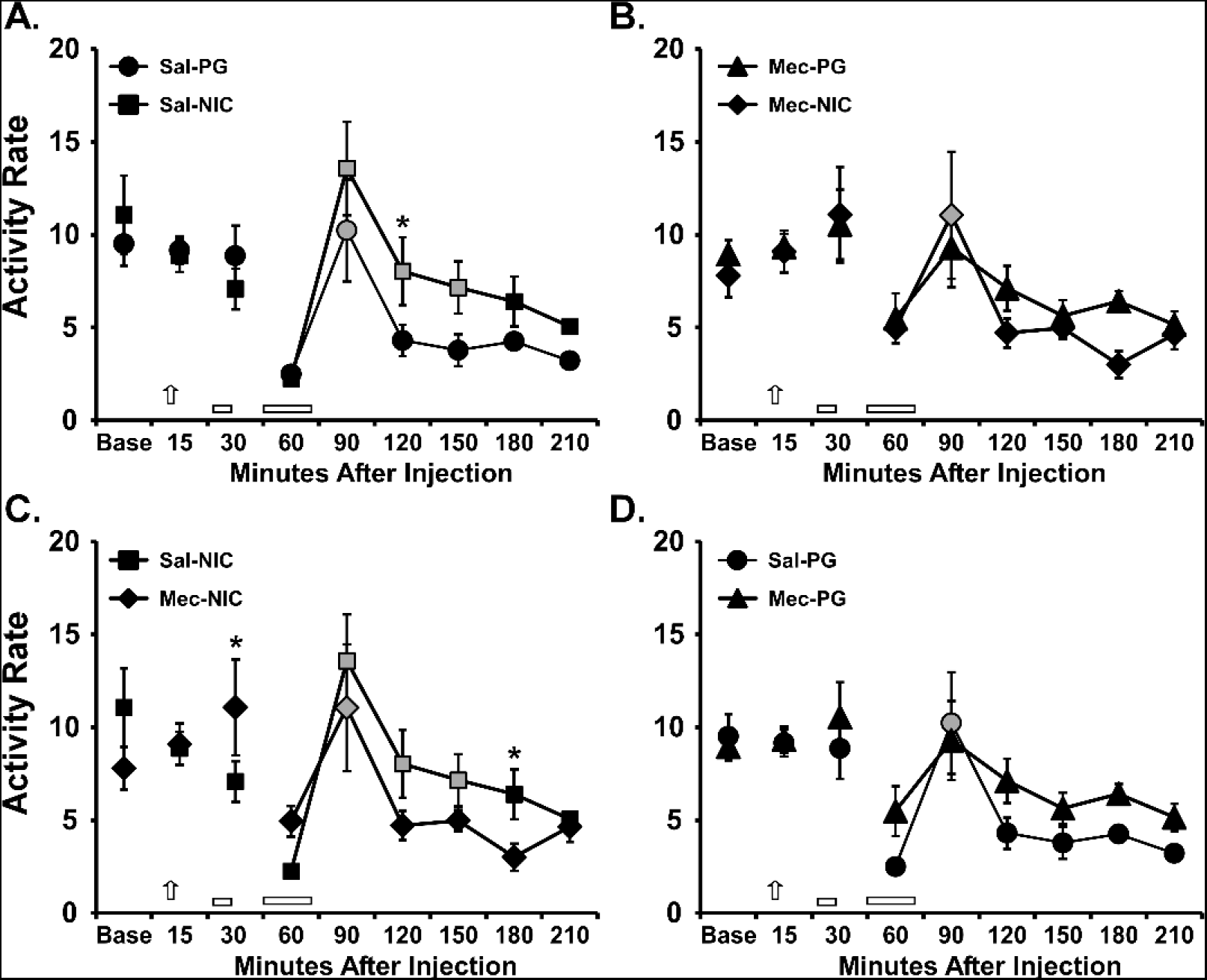
Mean (±SEM; N=8 male) activity rate after inhalation of the PG vehicle for 15 minutes and then either PG or nicotine (NIC; 30 mg/mL; 15 min) for 30 minutes one hour later. The four conditions were analyzed together but are presented in pairs for clarity. The session time is relative to the initial saline/mecamylamine injection. Arrows indicate timing of pre-session injection of <u>Sal</u>ine or <u>Mec</u>amylamine and the bars indicate the inhalation intervals. Shaded symbols indicate a significant difference from the 60-minute timepoint, within treatment condition. For a given time-point, a significant difference between inhalation conditions is indicated by ^*^.

### Effect of subcutaneous nicotine doses (0.1-0.8 mg/kg) and mecamylamine (2 mg/kg) in adult female rats

The activity of **<u>Group 2</u>** female rats was increased by NIC injection (**Figure 8A**), and the ANOVA confirmed significant effects of Time Post-injection [F (7, 49) = 19.51; P < 0.0001], and of the interaction of Time and Dose [F (28, 196) = 1.70; P < 0.05]. Activity was significantly different from the baseline 30 minutes after injection of NIC 0.1-0.8 mg/kg, but not after the saline injection. Activity did not differ compared with saline after the 0.1 mg/kg dose, but the post-hoc test confirmed significant differences from saline after 0.2 and 0.4 mg/kg (30 minutes) or 0.8 mg/kg (30 and 120 minutes) doses. The *temperature* of female rats was significantly affected by Time Post-injection [F (7, 49) = 6.01; P < 0.0001], and by the interaction of Time with Dose [F (28, 196) = 3.81; P < 0.0001] (**Figure 8C**). The hyperactivity of female rats induced by NIC, was attenuated by administration of mecamylamine (**Figure 8B**). The ANOVA confirmed significant effects of Time Post-injection [F (8, 56) = 7.54; P< 0.0001], and of dose Conditions [F (3, 21) = 3.31; P < 0.05], but not of the interaction of factors. Activity was significantly different from the baseline after injection of the 0.8 mg/kg (30 minutes) dose, but not in the saline, mecamylamine and mecamylamine-NIC dosing conditions. Activity did not differ compared with saline in the 0.8 mg/kg NIC, 2 mg/kg mecamylamine and their combination condition, but the post-hoc test confirmed significant differences in activity 30 minutes after the 0.8 mg/kg NIC compared with the mecamylamine condition. The *temperature* of female rats was not significantly affected by Time Post-injection, but significant effects of the Dose Condition [F (3, 18)= 7.07; P < 0.01] and of the interaction of Time and Dose [F (24, 144) = 2.31; P < 0.01] were confirmed (**Figure 8D**).

**Figure 8.**
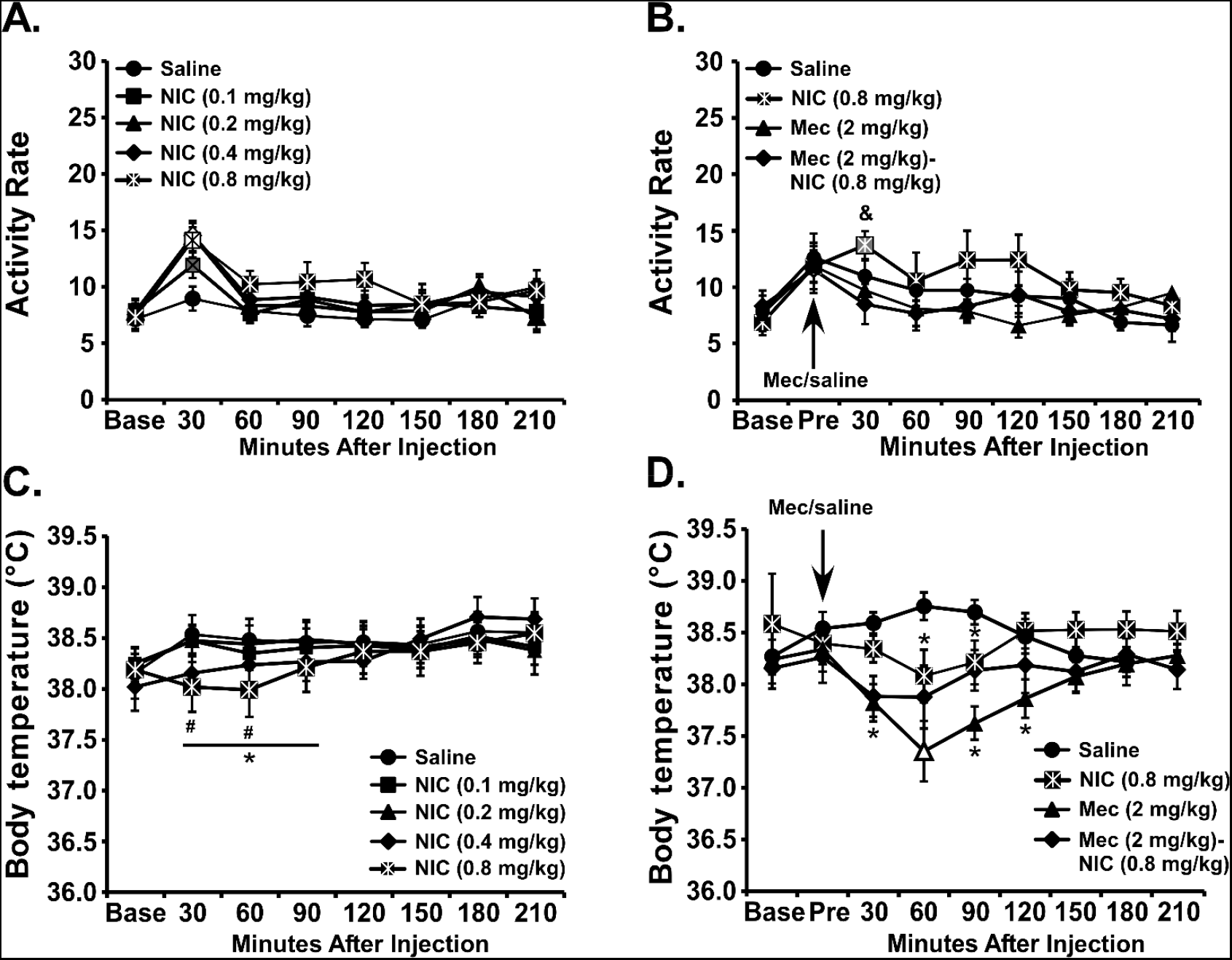
Mean (±SEM; N=8 female) A, B) activity rate and C, D) temperature after injection of the vehicle or nicotine (NIC; 0.1, 0.2, 0.4 and 0.8 mg/kg). Open symbols indicate a significant difference from both the Saline condition at a given time-point and the within-treatment baseline, while shaded symbols indicate a significant difference from the baseline, within treatment condition, only. A significant difference of NIC (0.8 mg/kg) from saline is indicated by ^*^ and a significant difference of NIC (0.4 mg/kg) from saline is indicated by ^#^. A significant difference from mecamylamine across NIC condition is indicated with &.

### Effect of subcutaneous nicotine doses (0.1-0.8 mg/kg) in adult male rats

The *activity* of the **<u>Group 2</u>** male rats was generally increased by the injection (**Figure 9A**) and the ANOVA confirmed significant effects of Time Post-injection [F (6, 42) = 25.54; P<0.0001], and of the interaction of Time with Dose Condition [F (24, 168) = 1.98; P<0.01]. The post-hoc test confirmed that activity was significantly different from the baseline at the 30-minute timepoint for all conditions. Although differences were small, the post-hoc test further confirmed that activity was higher after the 0.2 and 0.8 doses compared with saline and higher after the 0.2 dose compared with the 0.1 dose (see **Figure 9A inset**). The *temperature* of male rats was significantly affected by Time Post-injection [F (6, 42) = 3.67; P<0.01], but not by Dose Condition or by the interaction of factors (**Figure 9C**). The post-hoc test confirmed that across Pre-treatment conditions, temperature was elevated 180 minutes after injection compared with the baseline as well as 60- and 90-minutes post-injection.

**Figure 9.**
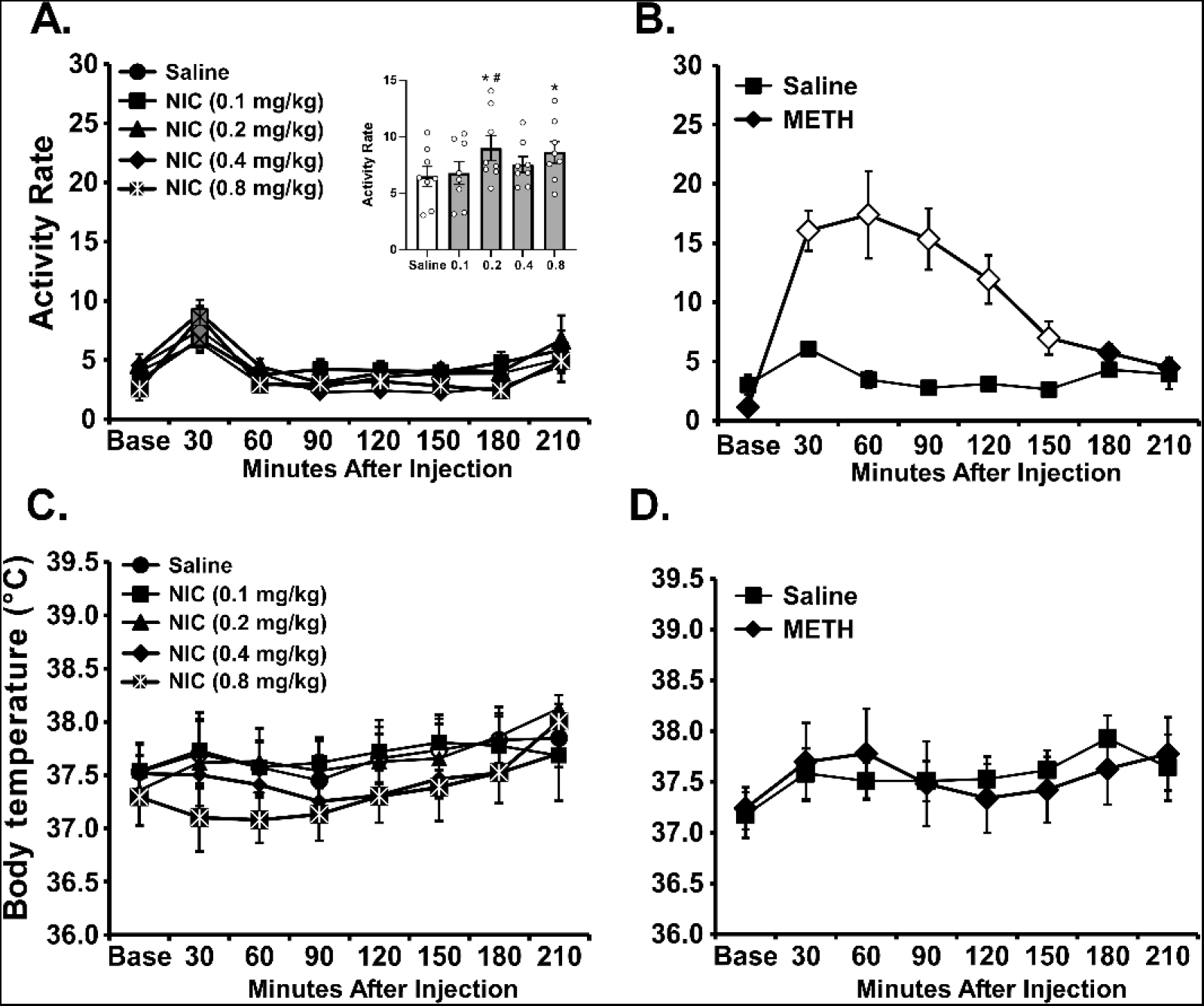
Mean (±SEM; N=8 male) A, B) activity rate and C, D) temperature after injection of A, C) nicotine (NIC; 0.0, 0.1, 0.2, 0.4 and 0.8 mg/kg, s.c.) or B, D) methamphetamine (0.0, 1.0 mg/kg, s.c.). The inset in A depicts the entire session activity average. Open symbols indicate a significant difference from both the baseline, within treatment condition, and saline at the respective timepoint. A significant difference from saline is indicated by ^*^ and from the 0.1 dose with # in the inset.

As a positive control it was found that the activity of male rats was increased by METH injection (**Figure 9B**); the ANOVA confirmed significant effects of Time Post-injection [F (6, 42) = 14.90; P<0.0001], of Pre-Treatment Condition [F (1, 7) = 23.87; P<0.005] and of the interaction of factors [F (6, 42) = 15.17; P<0.0001]. The post-hoc test confirmed activity was significantly different from the baseline 30-150 minutes after 1 mg/kg METH injection, but not after the saline injection. The post-hoc test further confirmed activity was higher from 30-150 minutes after METH compared with the same timepoints after saline injection. There was only a significant effect of Time Post-injection [F (6, 42) = 4.73; P<0.001] on body temperature and the post-hoc test confirmed that across Pre-treatment conditions, temperature was elevated 30-60 and 180 minutes after injection compared with the baseline (**Figure 9D**).

### Plasma nicotine and cotinine after s.c. and vapor inhalation

Nicotine produced dose-dependent effects on plasma nicotine and cotinine in **Group 4** male Wistar rats **(Figure 10A)**. The ANOVA confirmed significant effects of nicotine dose [F (2, 14) = 62.83; P<0.0001], of analyte [F (1, 7) = 118.0; P<0.0001], and of the interaction of factors [F (2, 14) = 29.03; P<0.0001]. The post-hoc test confirmed a significant difference after the 0.8 mg/kg dose compared with the 0.2 and 0.4 mg/kg doses, for both plasma nicotine and cotinine, as well as between the 0.2 and 0.4 mg/kg doses for nicotine. Nicotine concentration was significantly higher than cotinine concentration after all doses.

**Figure 10.**
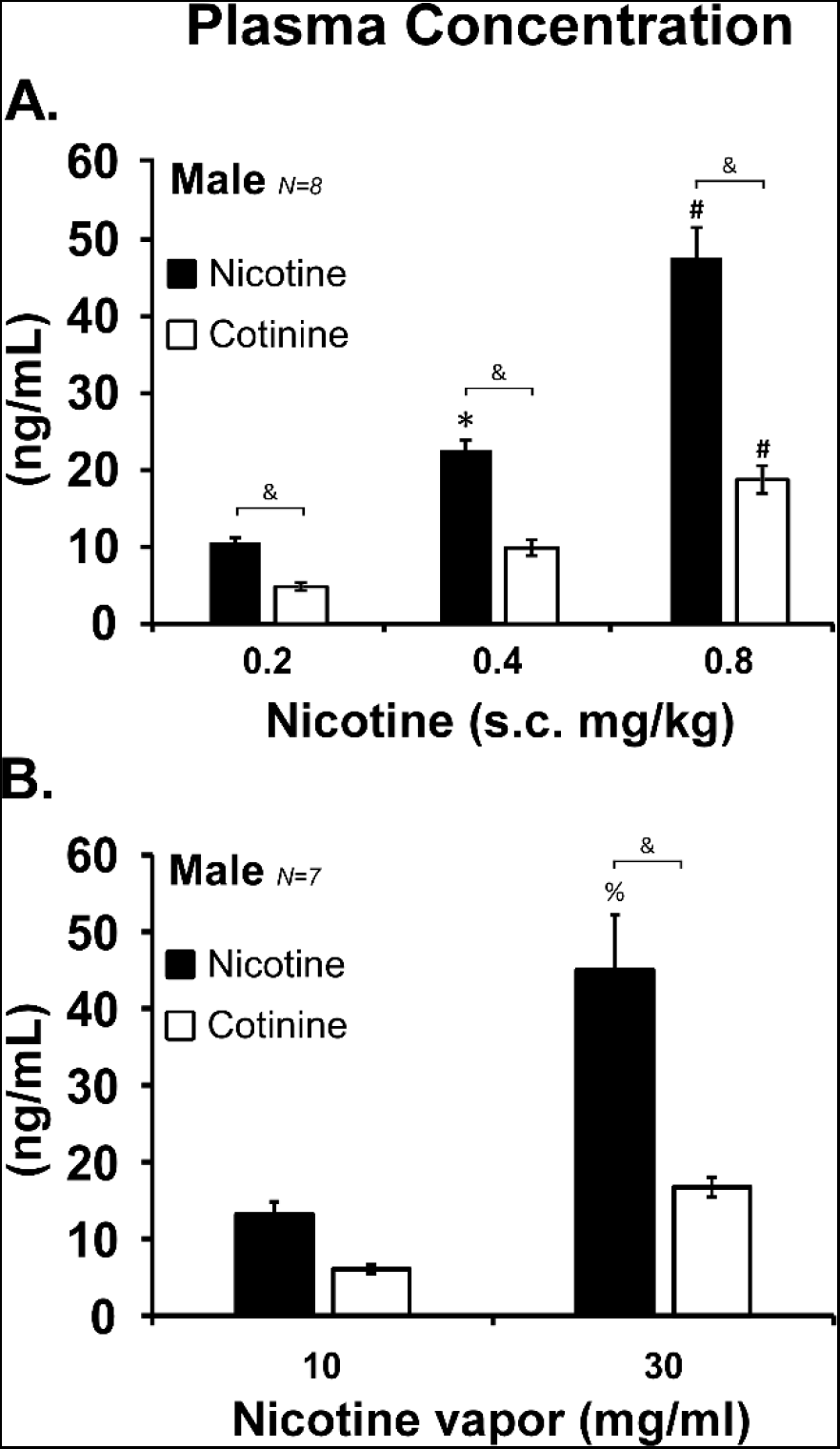
Mean (±SEM) plasma nicotine and cotinine in male Wistar rats after A) injection of nicotine (0.2-0.8 mg/kg, s.c.) 15 minutes before blood collection and B) vapor inhalation of nicotine (10 and 30 mg/ml in the PG vehicle) for 30 minutes. A significant difference from all other s.c. doses within analyte is indicated with #, a difference from the 0.2 mg/kg s.c. dose with ^*^, and a significant difference from the 10 mg/ml inhalation condition with %. A significant difference between nicotine and cotinine concentrations is indicated with &.

The inhalation of nicotine also produced dose-dependent effects on plasma nicotine and cotinine in the **Group 4** male rats **(Figure 10B)**. The ANOVA confirmed significant effects of nicotine concentration [F (1, 6) = 19.69; P<0.005], of analyte [F (1, 6) = 40.59; P<0.001], and of the interaction of factors [F (1, 6) = 9.91; P<0.05]. The post-hoc test confirmed a significant difference between nicotine and cotinine levels after the 30 mg/mL concentration, and a significant difference between 10 and 30 mg/mL concentrations for plasma nicotine, not cotinine.

## Discussion

The present study confirms that nicotine inhalation increases the locomotor activity of male and female Wistar rats, albeit to a larger extent in female rats under ∼equivalent dosing conditions. Inhalation of vapor from a 30 mg/mL concentration of nicotine in the vehicle was effective, consistent with our previous findings in male Sprague-Dawley rats (Javadi-Paydar et al. 2018). Nicotine injection (0.1-0.8 mg/kg, s.c.) only minimally increased the activity of male rats and modestly increased the activity of female rats in this study, despite the fact that the tested inhalation and injection conditions produced overlapping plasma concentrations of nicotine and cotinine in both male and female Wistar rats. Nicotine-induced hyperactivity depended on dose when altered by means of exposure duration in male rats (**Figure 2**) and by changing the drugconcentration in the vapor vehicle in female rats (**Figure 4**). Locomotor effects of nicotine inhalation were prevented by mecamylamine administration in female rats (**Figure 5**), and to some extent in male rats (**Figure 7**), thereby confirming the pharmacological specificity of the effect. Thus, the data show that the EDDS approach is successful in delivering behaviorally relevant doses to both male and female Wistar rats and may alter locomotor activity more consistently than when nicotine is injected. There was consistently a smaller effect of nicotine on locomotor activity in male rats compared with the response in female rats. This was true when contrasting effects of a nicotine (30 mg/mL) inhalation for 30 minutes in Group 1 rats (**Figure 1 A, B**), effects of a nicotine (30 mg/mL) inhalation for 15 minutes in Group 2 rats (**Figure 4A** vs **Figure 6A**), or the impact of subcutaneous injection of nicotine in Group 2 rats (**Figure 7A** vs **Figure 8A**). This sex difference is similar to the impact of s.c. nicotine injection on open field locomotor activity in male and female Sprague-Dawley rats (Gutierrez et al. 2024b). In the current inhalation studies, male rats often expressed an increase in activity after PG inhalation (**Figures 1B, 6A**) which obscured any additional effect of nicotine. This contrasted with the female rats, which expressed a higher locomotor response to nicotine and less response to vehicle vapor. Although the hyperlocomotor response in male rats was most robustly different from the PG condition after a 40-minute single nicotine inhalation session, the Group 1 males also exhibited a habituated PG response in that study which unmasked an effect of the 30-minute inhalation (**Figure 2**). As in our prior study (Javadi-Paydar et al. 2018), evaluating multiple vapor episodes in a single session also habituated male rats to the PG response and unmasked a drug effect (**Figure 6A**). Nevertheless, nicotine produced only modest locomotor stimulant effects in male rats, even following 0.2-0.8 mg/kg, s.c. injection. This wasn’t because the male rats of that age couldn’t express elevated locomotor activity, since robust locomotor stimulation was observed after an injection of methamphetamine (**Figure 9B**).

Hypothermic responses to nicotine were observed after a single 30-minute inhalation exposure or a 0.8 mg/kg injection in female rats (**Figures 1C, 8C**), but this was not observed in all studies in females, nor in male rats in this study. Nicotine inhalation reduced body temperature in male and female peri-adolescent and adult male Sprague-Dawley rats (Gutierrez et al. 2024b; Javadi-Paydar et al. 2019b). A similar hypothermic effect of nicotine was reported 10 min after s.c. injection in adolescent and adult female Sprague-Dawley rats in a prior study (Levin et al. 2003). This apparent strain difference is similar to a comparable insensitivity of Wistar rats to the hypothermic effects of THC (Taffe et al. 2021). Interestingly, Wistar rats are more sensitive than Sprague-Dawley rats to the hypothermic effects of 4-methylmethcathinone, and similarly responsive to hypothermia induced by 8-OH-DPAT (Wright et al. 2012), suggesting a complex interaction of strain with multiple temperature disrupting mechanisms. Although the *locomotor* impact of nicotine was blocked by mecamylamine in the current study, the hypothermic response to nicotine inhalation in female rats (**Figures 5C**) and to nicotine injection in male rats (**Figure 8D**) was *increased* by mecamylamine. This is likely because mecamylamine itself reduced body temperature in the absence of nicotine here and in prior studies (Hetzler and Bauer 2013). The additive effect on lowering body temperature has also been reported in mice treated with mecamylamine prior to nicotine vapor inhalation (Honeycutt et al. 2020).

Plasma levels of nicotine in these female rats were higher than those in males after inhalation (**Figure 3**), which is consistent with studies which showed female rats have higher plasma and brain nicotine levels after injection (Harrod et al. 2007; Rosecrans 1972) and after inhalation in our prior report (Gutierrez et al. 2024b). It cannot be ruled out that sex differences in locomotion after inhalation may be due to dose differences between male and female rats. However, the mean sex difference after inhalation was minor, and furthermore, the males exhibited slightly *higher* plasma nicotine levels after injection and yet showed no locomotor impact of nicotine injection. Plasma nicotine levels varied in male rats after 0.2 vs 0.8 mg/kg injection to a larger extent (**Figure 10A**) than any observed sex differences, and yet the female rats exhibited a similar degree of locomotor stimulation across this dose range (**Figure 8A**). Thus, the most likely interpretation is that female rats react more strongly than do males to the locomotor effects of nicotine.

Although female rats may have a 3-to 5-fold slower nicotine metabolism rate than that of males, leading to longer nicotine half-life and lower plasma cotinine levels (Kyerematen et al. 1988; Rosecrans 1972; Sanchez et al. 2014; Torres et al. 2009), there was no evidence of different cotinine levels at the limited timepoints assessed in this study (**Figures 3**). The cotinine levels in males and females after a 30-minute inhalation session were also similar to the levels reported after 1 hour of nicotine vapor self-administration in male and female Wistar rats (Lallai et al. 2021), which underlines the relevance of our non-contingent dosing conditions. However, considerably higher nicotine levels were reported after voluntary nicotine vapor exposure in male and female Wistar rats in another study (Smith et al. 2020), thus future investigations would be valuable to resolve apparent differences across laboratories and inhalation approaches.

In summary, this study confirmed that vapor inhalation of nicotine using the EDDS approach results in locomotor stimulation in both male and female Wistar rats. Locomotor stimulant effects were less robust in male rats, but this was partially attributable to a stronger locomotor response to vehicle vapor. The magnitude of effect depended on dose, as manipulated by either length of inhalation session or the nicotine concentration in the vapor vehicle and the effects were attenuated by pre-treatment with the antagonist mecamylamine; together this demonstrated pharmacological specificity. These locomotor stimulant effects also occurred under dosing conditions that generated plasma nicotine levels overlapping with those produced by subcutaneous nicotine injection in behaviorally relevant doses.

## Acknowledgements

The study was conducted with the support of USPHS grants (R01 DA035281, R01 DA042211) and the Tobacco Related-Disease Research Program (TRDRP; T31IP1832). The NIH/NIDA and the TRDRP had no role in study design, collection, analysis and interpretation of data, in the writing of the report, or in the decision to submit the paper for publication. The authors are grateful to Eric L. Harvey, Ph.D., for contributions to the assessment of plasma nicotine and cotinine. The authors are grateful to Maury Cole and La Jolla Alcohol Research, Inc. for contributions to developing the vapor inhalation equipment and methods used in this study.

